# Genetic rescue of disrupted synaptic protein interaction network dynamics following *SYNGAP1* reactivation

**DOI:** 10.64898/2025.12.02.691969

**Authors:** Vera Stamenkovic, Felicia Harsh, Breann Kniffen, Gavin Rumbaugh, Stephen Smith

**Affiliations:** Center for Integrative Brain Research, Seattle Children’s Research Institute, Seattle, WA, USA; Department of Neuroscience, The Herbert Wertheim UF Scripps Institute for Biomedical Innovation & Technology, Jupiter, FL, USA; Department of Pediatrics, University of Washington, Seattle, WA, USA; Graduate Program in Neuroscience, University of Washington, Seattle, WA, USA

## Abstract

Synaptic protein interaction networks (PINs) dynamically translate neural activity into biochemical signals that regulate synaptic structure and plasticity. Disruption of these coordinated networks is a common feature of autism spectrum disorder (ASD) risk genes, yet it remains unclear whether the molecular organization of a perturbed network can be restored after development. Here, we examined how post-developmental re-expression of the synaptic Ras GTPase-activating protein SynGAP1 affects network structure and signaling dynamics in a conditional SynGAP1 haploinsufficient mouse. Quantitative multiplex co-immunoprecipitation (QMI) across development revealed that SynGAP haploinsufficiency selectively reduced SynGAP-containing complexes without broadly disrupting NMDA-dependent network responses. Tamoxifen-inducible re-expression of SynGAP at postnatal day 21 fully restored both steady-state and activity-dependent interactions within the SynGAP module in hippocampus, and additionally normalized secondary alterations in Shank–Homer scaffolding complexes in somatosensory cortex. These data demonstrate that biochemical restoration of a disrupted synaptic network is achievable, even after early developmental windows have closed. Our findings suggest that while critical periods may constrain functional recovery, molecular network normalization remains possible through genetic reactivation of haploinsufficient synaptic regulators.

## Introduction

Protein–protein interactions form molecular logic circuits that enable cells to interpret inputs and transition between homeostatic states^1–3^. The dynamic activity of protein interaction networks (PINs) is crucial for translating external stimuli into intracellular responses, ultimately orchestrating complex processes such as growth, migration, or synaptic plasticity. At the glutamate synapse, a coordinated PIN comprised of neurotransmitter receptors, scaffolds, and effectors including kinases, phosphatases and ubiquitin ligases responds to neurotransmitter release by reorganizing its molecular associations^4–7^. In the resting state, the network stabilizes around a defined homeostatic setpoint; following depolarization, neurotransmitter agonists^5^, homeostatic scaling^8^, or behavioral learning^9^, post-translational modifications alter protein binding affinities and remodel the network’s topology. These changes concurrently reshape receptor scaffolding to adjust synaptic strength, and activate kinase cascades that propagate intracellular signals. Thus, the configuration of a PIN both orchestrates the cell’s functional response to an incoming input, and encodes its recent sensory experience.

Autism-linked mutations cluster into functional PINs that are important for neuronal activity-dependent plasticity^10,11^. Analyses of autism risk gene function highlight a distributed pathway responsible for neuronal activity-dependent plasticity and homeostasis, with mutations clustering into networks involving synaptic/dendritic function^12^ (represented by ASD risk genes including *SHANK3, SYNGAP1, SCN2A, ANK2* ^13,14^), activity-dependent DNA translational regulation (*CHD8*, *ARID1B*, Baff complex ^15^), and activity-dependent mRNA translation (*FMR1*, *PTEN*, *PIK3CA*, mTOR signaling^16^). These findings support the view that autism arises from disruption of neuronal homeostatic control systems that act across developmental time to constrain synaptic activity and coordinate the expression of species-appropriate social behavior^12^. The distributed and interconnected nature of these networks means that individual gene mutations can propagate through multiple molecular pathways, altering both acute PIN responses to stimuli and the long-term setpoints around which synaptic signaling equilibrates^17^.

As the field begins to understand molecular mechanisms and test treatments to restore the function of disrupted ASD risk genes, it is unclear how established homeostatic adaptations within PINs will respond to late interventions. It is widely appreciated that developmental critical periods exist in brain circuits; once a given circuit has matured, restoration of wild-type-like molecular function can no longer normalize cellular or behavioral outcomes. For example, adult re-expression of Shank3 in conditional mutant mice normalized synaptic protein composition and social behavior, but failed to rescue repetitive behaviors or anxiety^18^. In Ube3a knockout animals, motor deficits can be rescued in adulthood, but anxiety, repetitive behavior and seizures were only rescued by early reactivation^19^. Clearly, rescue of complex behavior, particularly anxiety-like and repetitive behavior, is limited by circuit-level critical periods.

At the biochemical level, emerging evidence suggests that synaptic signaling networks themselves may become developmentally entrenched. In prior work from our group using Fmr1 knockout mice, metabotropic glutamate receptor antagonists or Src-family kinase inhibitors, which correct excessive protein synthesis and rescue behavioral deficits in adult animals, failed to restore wild-type patterns of synaptic protein-protein interactions^20^. Rather, drug treatment led to a stable, alternate network configuration involving increased Shank3 and NMDA receptor interactions not seen in wildtype animals. These data raise the possibility that once signaling networks have re-equilibrated to a mutant homeostatic state, later genetic or pharmacologic interventions may improve mouse behavioral function, but cannot fully restore the original PIN state. Here, through genetic re-expression of mouse *Syngap1* ^21–23^, we directly test whether post-developmental restoration of a member of the glutamate synapse protein interaction network can restore protein network function, or alternatively, if altered network homeostasis leads to continued abnormalities despite an ideal genetic rescue.

Rescue of *Syngap1* haploinsufficiency is a promising approach to test this hypothesis. First, rare pathogenic variants in *SYNGAP1* leading to genetic haploinsufficiency cause a neurodevelopmental disorder characterized by intellectual disability, epilepsy and autism^11,24–27^, demonstrating that that reduced SynGAP protein disrupts human cognition, neural network excitability, and adaptive behavior. Second, in animal models, reduced SynGAP protein abundance caused by *Syngap1* genetic haploinsufficiency results in well described disruptions to synapse structure and function^22,28^. These alterations arise due to its crucial functions within the postsynaptic density; SynGAP is a synaptic Ras GTPase-activating protein that undergoes liquid-liquid phase separation with PSD-95, forming synaptic condensates that organize AMPA receptor nanodomains and influence postsynaptic receptor clustering^6^. Following depolarization, SYNGAP is phosphorylated by CaMKII in an NMDA receptor-dependent manner, leading to its rapid dispersion from the PSD and the opening of binding slots for AMPA receptor recruitment^29,30^. Dynamic SynGAP protein-protein interactions regulate synaptic strength and cognitive functions independently of SynGAP’s enzymatic activity by competing with AMPAR-TARP complexes^31^, highlighting an increasing appreciation of the direct role of PIN dynamics (rather than enzymatic activity) in synaptic plasticity^32,33^ (but see ^34^). Non-synaptic developmental changes have also been reported^35^. Third, a *Syngap1*^+/ls^ mouse has been developed in which a lox-stop-lox cassette causes germline *Syngap1* haploinsufficiency that is genetically reversable using tamoxifen-Cre^22^. Using this model, it has been shown that re-expression of deficient SynGAP protein in mice caused by genetic haploinsufficiency after developmental critical periods (e.g., >PND60) rescues deficient long-term potentiation^36^ and reactivates experience-dependent sensory-evoked ensemble plasticity^37^. This approach also boosts the power of hippocampal theta rhythms, improves deficient memory consolidation, and mitigates a variety of seizure-related functional measures^23,38^. Consistent with findings from other rare variant gene reactivation models, re-expression at post-developmental timepoints fails to rescue some behavioral and cellular features of the model^21,22^, most likely due to persistent hard-wired disruptions to developmental functional network connectivity as demonstrated through various cell-specific/region specific rescue strategies^39,40^. Despite the mixed functional rescue results in behavior, there is substantial evidence that restoration of low SynGAP abundance in the *Syngap1*^+/ls^ system rescues synapse- and circuit-level plasticity. Thus, this model provides an ideal substrate to test if normalization is possible at the molecular level, or if developmental changes to PIN structure are irreversible.

We report that SynGAP haploinsufficiency produces alterations to the synaptic protein network that are largely confined to SynGAP-containing interactions. Re-expression of SynGAP after PND21 rescues both steady-state and activity-dependent SynGAP interactions, and normalizes the protein network to a wild-type-like state. To our knowledge, this is the first example of a synaptic biochemical network being normalized postnatally by genetic re-expression, and strengthens the premise for treatment strategies aimed at genetic rescue of haploinsufficiency^38,41^.

## Results

### SynGAP haploinsufficiency alters hippocampal protein interaction networks

To observe the effect of SynGAP haploinsufficiency on dynamic synaptic protein interaction networks, we treated acute hippocampal slices from P60 SynGAP^+/ls^ mice with NMDA/Glycine to mimic strong Ca^2+^ flux associated with synaptic potentiation, or with DHPG to induce weaker metabotropic signaling associated with synaptic depression, as previously described^5,20^. Following brief (5min) treatment, quantitative multiplex co-immunoprecipitation^42,43^ was used to monitor 400 binary interactions among 21 proteins critical for synaptic signal transduction (antibodies in Table S1). Correlation network analysis^42,44^, which clusters protein-protein interactions based on correlated behavior across N=24 samples (Fig 1A,B), identified four modules of co-regulated interactions. One module correlated most strongly with NMDA treatment (correlation coefficient (CC) = −0.92, p = 3×10^-10^), and one module with genotype (CC = −0.59, p = 0.002) (Fig 1C). A heatmap (Fig 1D) and node-edge diagram (Fig 1E,F) display interactions that are both a member of one of these two modules, and are significantly different between treatment groups by ANC, an adaptive nonparametric statistical test corrected for multiple comparisons that was designed specifically for QMI data^42^.

**Figure 1:**
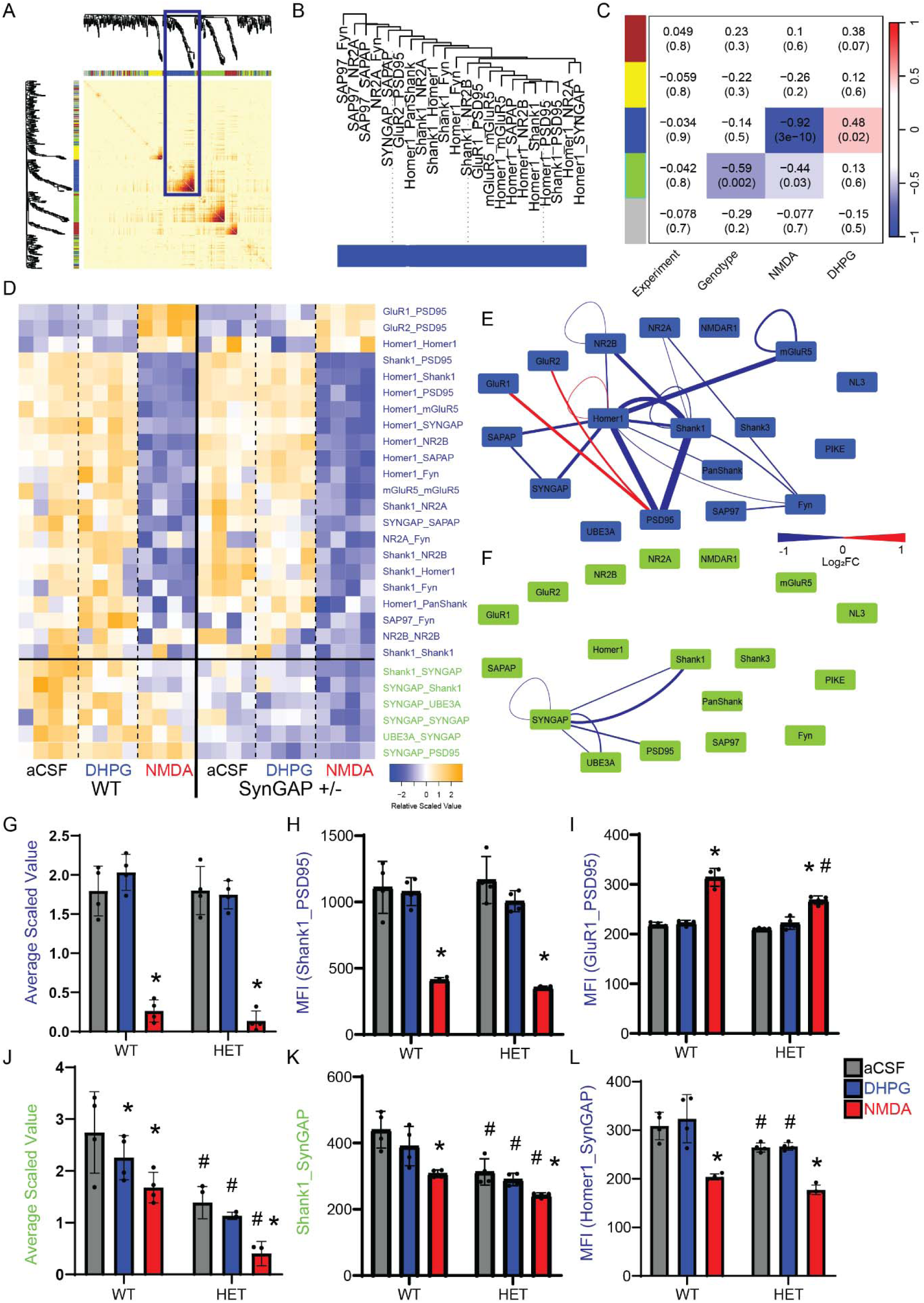
Protein interaction network response to NMDA in P60 SynGAP^+/ls^ hippocampal slices. A) Toplogical overap matrix shows clustering of individual interactions (dendrogram tips) into modules. Red color indicates more correlated intractions. B) An example of protein interactions clustered in the “blue” NMDA-responsive module. Note the diversity of IP and probe antibodies showing correlated behavior. C) A module-trait table shows the correlation coeficent (top number) and p-value (bottom number) for the correlation between each module’s eigenvector and experimental variables listed at bottom. D) A row-scaled heatmap shows all interactions that were both significant in binary comparisons by adaptive nonparametric test corrected for multiple comparisons (ANC) and a member of a CNA module. Interactions listed at right are colored by module membership. E,F) Node-edge diagrams show interactions in the NMDA (E) and SynGAP (F) modules. Edge width indicates the magnitude of the fold-change for NMDA vs aCSF or WT vs SynGAP^+/ls^, and edge color indicates an increase or decrease. G-L) Bar graphs represent the average scaled value (G,J) and example interactions (H-I, K-L) for the blue NMDA-correlated (G-I, L) or the green SynGAP-correlated (J-K) modules. * indicates p<0.05 comparing aCSF vs treatment within-genotype; # indicates p<0.05 comparing WT vs. SynGAP^+/ls^ within-treatment, by 2-way ANOVA followed by Tukey’s post-hoc testing.

The NMDA-correlated module, arbitrarily color-coded blue, consists of 22 interactions that almost all dissociate upon NMDA stimulation. The average scaled value of the module, which represents the collective behavior of its interactions and may be more robust than measurements of single interactions^8^, was significantly reduced by NMDA stimulation, and was not different between WT and SynGAP^+/ls^ slices, indicating a grossly normal response to NMDA stimulation (Fig 1G). The interaction most strongly correlated with module behavior, IP Shank1 probe PSD95 (abbreviated Shank1_PSD95), was reduced more than 50% by NMDA stimulation in both genotypes (Fig 1H). GluR1_PSD95 (Fig 1I) and GluR2_PSD95 increased following NMDA treatment, and were increased significantly less in SynGAP^+/ls^ slices. These data are consistent with current models of synaptic plasticity leading to the dissociation of synaptic scaffolds and SynGAP^29^, which opens binding slots for recruitment of new AMPA receptors^30^; in SynGAP^+/ls^ mice, lower levels of SynGAP mean that fewer binding slots are opened, and fewer AMPA receptors are recruited. However, besides this relatively minor difference, the network response to NMDA remained largely intact in SynGAP^+/ls^ hippocampi.

The average scaled value of the genotype-associated module was reduced by both SynGAP heterozygosity and by NMDA treatment (Fig 1J), again consistent with activity-dependent SynGAP dissociation. The interaction most strongly correlated to the module, Shank1_SynGAP, was significantly reduced by NMDA in both WT and SynGAP slices, but the basal levels in SynGAP slices was almost 50% less than wildtype (Fig 1K). Interactions including SynGAP_SynGAP, and SynGAP_Shank1, _PSD95 and _UBE3A followed this trend. The dissociation of Homer1_SynGAP (Fig 1L) and SynGAP_SAPAP was so strong that CNA placed it in the NMDA module, but gentoype effects are still apparent. Overall, the data reveal that in slices from adult SynGAP^+/ls^ heterozygotes, levels of SynGAP are reduced, but protein complex dynamics in response to NMDA remain intact and grossly normal, with the exception of lower PSD95_GluR1/2 recruitment.

### Protein interaction network structure evolves across development

SynGAP1 expression is low during prenatal development, then peaks around P14 and declines to lower steady-state levels in adults^22^. To explore the developmental trajectory of SynGAP haploinsufficiency, we repeated the slice stimulation paradigm at P7 (pre-peak) and P14 (peak SynGAP). At both ages, we identified two modules of interest: one that correlated most strongly with NMDA (P7: CC = 0.82, p = 2×10^-8^, P14: CC = −0.93, p = 8×10^-11^) and one module that correlated most strongly with SynGAP haploinsufficiency (P7: CC = −0.74, p = 3×10^-6^, P14: CC = −0.86, p = 8×10^-8^) (Fig S1A and S2A). The NMDA module was notably weaker at P7, with fewer interactions reaching statistical significance, a lower magnitude of change of significant interactions, and a network dominated by interactions involving Homer1 (Fig S3 A-I). Low levels of Shank1 and PSD95 in early development (Fig S3K,L) resulted in interactions that were prominent at P60, like Shank1_PSD95 (Fig 1H), being absent at P7. GluR1_PSD95 was slightly but significantly increased by NMDA in P7 SynGAP^+/ls^ slices, not in WT (Fig S1E), consistent with early maturation of synaptic spines observed in SynGAP^+/ls^ cortex^21^. The P7 SynGAP module was not significantly reduced by NMDA (Fig S1F), although some interactions including SynGAP_SynGAP (Fig S1G) did show a reduction following NMDA treatment. Moreover, SynGAP_PSD95 was increased by NMDA in WT, and trended towards an increase in SynGAP^+/ls^ (Fig S1H).

By P14, the NMDA module contained 31 interactions (vs. only 13 at P7, Fig S2B), and the magnitude of change was greater, similar to adult slices (Fig S3D-F). NMDA stimulation resulted in strong dissociation of most module interactions in both genotypes (Fig S2C), while a few interactions, including GluR1_PSD95 (Fig S3E) was increased in both genotypes. The WT SynGAP module responded to both NMDA and DHPG in WT P14 hippocampi, and was consistently reduced in SynGAP^+/ls^ animals and failed to respond to NMDA (Fig S2F). SynGAP_SynGAP was consistently lower in SynGAP^+/ls^ slices, and while SynGAP_PSD95 increased with NMDA stimulation in WT slices, similar to P7, the magnitude of the interaction did not change with NMDA (Fig S3H).

Comparing across timepoints, many interactions increased consistently across development (P7<P14<P60), including GluR1_PSD95, PSD95_PSD95, and all interactions involving Shank1 (Fig S3H-J). Consistent with prior studies, SynGAP_SynGAP peaked at P14 (Fig S3K), although interactions containing SynGAP either plateaued at P14 (Homer1_SynGAP, Fig SL), or continued to increase into adulthood (PSD95_SynGAP, Fig SL), suggesting synaptic maturation. A few interactions such as Homer_GRM5 decreased with development (Fig S3N). Overall, these data demonstrate an evolving pattern of protein interaction network dynamics across development, with NMDA consistently producing dissociation among a number of synaptic protein complexes and an increase in PSD95_AMPA receptors that increased with age. SynGAP heterozygosity reduced SynGAP levels, and reduced the abundance of SynGAP in complexes that dissociated with NMDA. However, in contrast to Fragile X^20^ or Shank3^8^ mouse models previously described, the effects of SynGAP heterozygosity did not extend to non-SynGAP-containing complexes. Except for developmentally regulated changes in AMPA receptor recruitment to PSD95 (Fig S3J), the SynGAP+/- synaptic protein network responded normally to NMDA.

The response to DHPG, which produces smaller increases of intracellular calcium and weaker protein interaction changes compared to NMDA^9^, was weak in both genotypes, and few interactions reached statistical significance, preventing us from drawing conclusions about GRM5 signaling.

### Normalization of Protein networks by re-expression of SynGAP

Given the relatively limited scope of changes in SynGAP^+/ls^ hippocampal slices, involving almost exclusively SynGAP-containing complexes, SynGAP re-expression may offer an ideal proof-of-concept to test if PIN normalization is possible. We crossed SynGAP^+/ls^ mice to hemizygous mice expressing a tamoxifen (TMX)-inducible form of Cre recombinase. All generated genotypes, Cre^-^ and Cre^+^, WT and SynGAP^+/ls^ mice, were administered 75 mg/kg of TMX intraperitoneally for 3 consecutive days starting at P21 (Fig 2A), and the treatment was validated by Western blot when animals were 2 months old. Upon Cre activation, an artificial exon containing a premature stop codon is excised from the *SYNGAP1* gene^22^, leading to re-activation of the targeted allele and restoration of wild-type SYNGAP protein levels (Fig 2 B,C).

**Figure 2:**
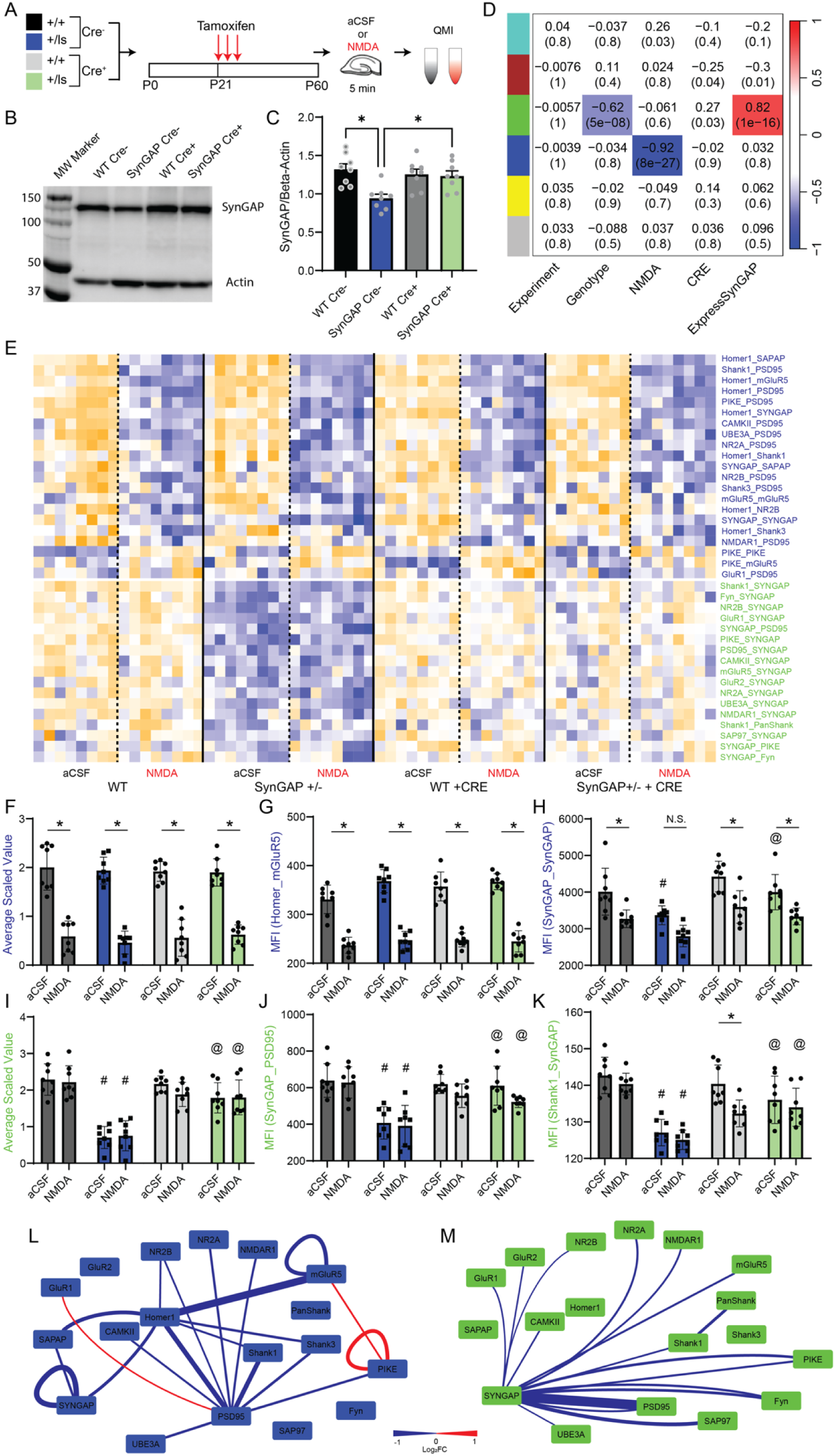
SynGAP re-expression in hippocampal slices. A) Experimental design B) Example western blot showing SynGAP levels at P60 across genotypes. C) Quantification of western blots. *p<0.05 by one-way ANOVA followed by Sidak post-hoc testing. D) A module-trait table shows the correlation coeficent (top number) and p-value (bottom number) for the correlation between each module’s eigenvector and experimental variables listed at bottom. E) A row-scaled heatmap shows all interactions that were both significant in binary comparisons by adaptive nonparametric test corrected for multiple comparisons (ANC) and a member of a CNA module. Interactions listed at right are colored by module membership. F-K) Bar graphs represent the average scaled value (F,I) and example interactions (G-H,J-K) for the blue NMDA-correlated (F-H) or the green SynGAP-correlated (I-K) modules. * indicates p<0.05 comparing aCSF vs treatment within-genotype/Cre; # indicates p<0.05 compared to WT/Cre^-^ within-treatment; @ indicates p<0.05 compared to SynGAP^+/ls^/Cre^-^ within-treatment, by 3-way ANOVA followed by Sidak’s post-hoc testing. L,M) Node-edge diagrams show interactions in the NMDA (L) and SynGAP (M) modules. Edge width indicates the magnitude of the fold-change for NMDA vs aCSF or WT vs SynGAP^+/ls^ respectively, and edge color indicates an increase or decrease.

Following P21 TMX, PND60 hippocampal slices were treated with NMDA or aCSF for 5 minutes, and CNA identified two modules of correlated interactions, one correlated with NMDA treatment (CC = −0.92, p = 8 x 10^-27^) and one correlated with SynGAP expression (CC = 0.82, p = 1 x 10^-16^) (Fig 2D). A heatmap of all significant interactions again showed a mainly dissociative network of interactions responding to NMDA, and interactions containing SynGAP protein reduced exclusively in SynGAP^+/ls^ Cre^-^mice (Fig 2E). The average scaled value of the NMDA module showed a significant reduction with NMDA (3-way ANVOA, F_1,36_ = 324.9, p < 0.0001), with no significant effect of genotype or Cre (Fig 2F). The interaction that showed the largest magnitude of change, Homer1_mGlurR5, was significantly reduced by NMDA in all conditions (Fig 2G). SynGAP_SynGAP clustered in the NMDA module because it was reduced following NMDA in WT and SynGAP^+/ls^ /Cre^+^ slices; it was not significantly reduced by NMDA in SynGAP^+/ls^ /Cre^-^, indicating a genotype effect consistent with prior experiments (Fig 2H). Moreover, the median fluorescent intensity (MFI) of SynGAP_SynGAP was significantly reduced in SynGAP^+/ls^ /Cre^-^ aCSF slices compared to WT, and was significantly increased comparing SynGAP^+/ls^ /Cre^-^ to SynGAP^+/ls^ /Cre^+^ slices in aCSF, confirming successful rescue of SynGAP protein levels, and of NMDA-dependent SynGAP dynamics (3-way ANOVA, NMDA F_1,56_= 49.26, p<0.001; CRE F_1,56_= 22.8, p<0.001; Genotype F_1,56_= 20.34, p<0.001;). Thus, while there were minimal differences in the NMDA module due to SynGAP heterozygosity, those that existed were normalized by TAM/CRE re-expression.

The average scaled value of the SynGAP module was reduced comparing WT with SynGAP^+/ls^ /Cre^-^, and increased comparing SynGAP^+/ls^ /Cre^-^ to SynGAP^+/ls^ /Cre^+^ (Fig 2I) (3-way ANOVA: CRE F_1,56_= 18.88, p<0.001; Genotype F_1,56_= 82.6, p<0.001; CRE X Genotype F_1,56_= 44.81, p<0.001). SynGAP_PSD95 (Fig 2J) and Shank1_SynGAP (Fig 2K) both showed significant rescue of basally reduced interactions. Node-edge diagrams showing the NMDA module (Fig 2L) and the SynGAP module (Fig 2M) highlight the contrast between extensive, network-wide changes induced by NMDA, and the more limited changes caused by SynGAP expression that all involve direct SynGAP complexes (except for a small change in Shank1_panShank). We conclude that SynGAP re-expression provided effective normalization of the glutamate synapse PIN in hippocampal slices.

### SynGAP in P60 mouse cortex

We next asked if SynGAP re-expression could also rescue protein interaction network deficits in the mouse primary somatosensory cortex. Following P21 re-expression in a new cohort of mice that was confirmed by western blotting (Fig 3A), S1 somatosensory cortical tissue was collected on P60 for QMI. CNA identified two modules that correlated with SynGAP haploinsufficiency, a SynGAP module (turquoise, cc = 0.77, p = 1 x 10^-16^) and a second “blue” module (cc = 0.6, p = 3 x 10^-9^) that was not identified in hippocampal tissue (Fig 3B). In contrast to the hippocampus, where the effects of SynGAP heterozygosity did not extend beyond SynGAP-containing complexes, the Blue module contained Shank/Homer complexes detected with non-SynGAP antibodies (Fig 3C), allowing us an opportunity to test if indirect effects of SynGAP loss would be normalized by re-expression.

**Figure 3:**
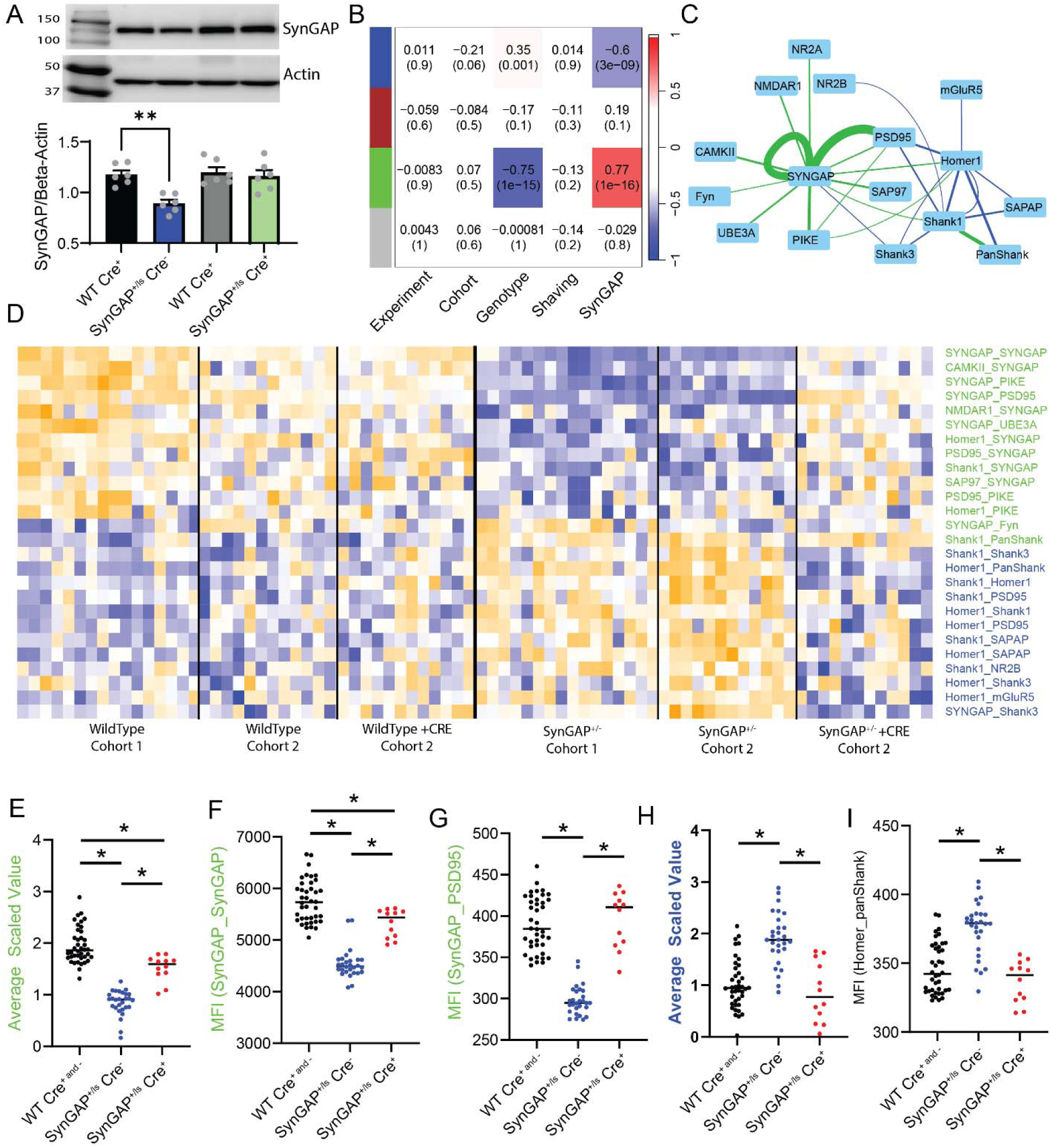
SynGAP re-expression in cortical slices. A) Example western blot showing SynGAP levels at P60 across genotypes, and quantification of western blots. *p<0.05 by one-way ANOVA followed by Sidak post-hoc testing. B) A module-trait table shows the correlation coeficent (top number) and p-value (bottom number) for the correlation between each module’s eigenvector and experimental variables listed at bottom. C) Node-edge diagram shows interactions in the SynGAP (green) and Shank/Homer (Blue) modules. Edge width indicates the magnitude of the fold-change for WT/Cre^-^ vs SynGAP^+/ls^/Cre^-^, and edge color indicates module membership. D) A row-scaled heatmap shows all interactions that were both significant in binary comparisons by adaptive nonparametric test corrected for multiple comparisons (ANC) and a member of a CNA module. Interactions listed at right are colored by module membership. E-I) Graphs represent the average scaled value (E,H) and example interactions (F-G, I) for the green SynGAP (E-G) or the blue Shank/Homer (H-I) modules. * indicates p<0.05 comparing for the selected comparison by one-way ANOVA followed by Tukey’s post-hoc testing.

The SynGAP module was significantly reduced in SynGAP^+/ls^ /Cre^-^ cortex, and rescued by TAM/Cre, although the average scaled value of the module remained below WT levels (ANOVA F_2,77_ = 112.7, p<0.0001), suggesting that TAM/CRE was less than 100% effective at restoring cortical SynGAP levels (Fig 3D,E). Indeed, SynGAP_SynGAP was significantly reduced in SynGAP^+/ls^ /Cre^-^, and while levels were higher in Cre^+^ vs Cre^-^ SynGAP^+/ls^ cortex, they were still significantly lower than in WT (Fig 3F). However, the levels of other SynGAP interactions, including SynGAP_PSD95 (Fig 3G), were normalized by SynGAP re-expression, similar to the hippocampus.

The average scaled value of the Homer/Shank module was increased in SynGAP^+/ls^ /Cre^-^ cortex, and reduced to WT levels following TAM/Cre (ANOVA F_2,77_ = 30.63, p<0.0001) (Fig 3H). The module was comprised of Homer1 and Shank1/3 interactions and included GRM5, SAPAP and PSD95 (Fig 3C). Interactions including Homer_panShank (Fig 3I) were increased in SynGAP^+/ls^ /Cre^-^ cortex and reduced back to WT levels in SynGAP^+/ls^ /Cre^+^ cortex. These data demonstrate that rescue of typical WT PIN composition and dynamics is possible as late as P21 in mouse cortical and hippocampal tissue.

## Discussion

Treatments aimed at correcting disease-associated biochemical abnormalities have met with limited success in neuropsychiatric disorders^45^. In addition to well-described critical periods in neuronal cell and circuit development limiting the potential treatment window to early stages of the lifespan, there is also the potential for biochemical pathways to become resistant to normalization. We previously showed, in the FMR1 knockout mouse model of Fragile X syndrome, treatments that normalized elevated protein synthesis and behavior failed to normalize protein interaction networks^20^. We documented widespread deficits in the synaptic PIN due to FMR1 mutation, then treated in vitro slices or mice with the GRM5 antagonist MPEP or the FYN kinase inhibitor Saracatinib. Both drugs were selected to counteract biochemical evidence of elevated GRM5 tone and increased FYN activity in the FMR1 mouse, and both drugs effectively normalized KO mouse phenotypes^20,46^. However, the drugs did not normalize PIN measurements, but rather induced a new, unexpected set of changes involving NMDA receptors and Shank scaffolds. The fact that the biochemical system did not return to a wild-type-like state, but instead settled on a third state that functionally improved behavior but resulted in novel biochemistry, begged the question, is normalization of a disrupted PIN even possible?

To approach this question, we used a genetic re-expression strategy to induce an optimal rescue of gene function^21–23^. Alternative strategies such as a viral vector delivery of a corrected gene or treatment with small molecule drugs would be more clinically relevant, but would risk incomplete rescue or off-target drug effects, confounding a potentially negative result. By removing a lox-stop-lox cassette and restoring somatic gene function, we were able to demonstrate that biochemical normalization of protein networks is possible.

SynGAP haploinsufficiency may be an unusually amenable phenotype to rescue, because the observed PINs effects were largely limited to its own interactions. In the hippocampus, we only detected a single non-SynGAP interaction in the genotype module, Shank1_panShank, which was also normalized by re-expression. In the cortex, we identified a greater number of Shank and Homer interactions that were upregulated in SynGAP, but were normalized by re-expression. The fact that the network changes were not as widespread compared to other models^8,47^ may be due to presence of a half-dose of SynGAP in the haploinsufficient state. Full knockout or gain-of-function deficits may be harder to normalize. Regardless, our result does provide encouragement that strategies to restore full gene expression in haploinsufficient models after development may normalize altered biochemistry^38,41^, as has been demonstrated for network- and behavioral-level substrates linked to hyperexcitability and memory consolidation^23^. Furthermore, these results also highlight the fundamental importance of the SynGAP-related PIN. While its network is relatively restricted, disrupting it has serious consequences for biochemical signaling at synapses^48^, while restoring its expression after development can reactivate synapse- and circuit-level plasticity that coincides with rescue of associated kinase signaling pathways^36,37^.

There are several limitations to this work. First, we were not able to show consistent activity-dependent PIN remodeling in response to DHPG in the hippocampus, as we have previously shown in the cortex^5,8,20^. This may be due to region-specific differences, or because the NMDA and genotype effects were so large as to statistically obscure the more subtle effect of DHPG. Similarly, whisker shaving failed to produce the expected activity-dependent PIN remodeling in the barrel cortex^8^, possibly due to inconsistencies in the dissections. Nevertheless, we were able to document genotype-specific, activity-modulated differences in PIN structure that were rescued by re-expression of SynGAP. Finally, our approach was limited to the biochemical effects of rescue, and did not examine known deficits in SynGAP mice such as synaptodendritic structure^21^ or behavior^22^ that have been previously documented to be resistant to rescue. So, while we have shown that molecular interaction deficits can be normalized, changes in neuronal morphology or circuitry that are known to be resistant to re-expression will limit future treatment approaches.

In conclusion, this study was narrowly designed to ask if biochemical rescue of PIN deficits in a genetic mouse model of autism were possible. The data indicate that re-expression of SynGAP results in complete normalization of the synaptic protein interaction network, including both SynGAP interactions and other downstream effects of haploinsufficiency involving the Shank-Homer scaffold. While critical periods of brain development will remain an important obstacle in long-term, durable treatment of neuropsychiatric disorders, this study suggests that biochemical normalization is within the realm of possibility.

## Materials and methods

### Animals

All work with animals was performed in compliance with the Seattle Children’s Research Institute Institutional Animal Care and Use Committee under approved protocol #00072 and federal guidelines. The generation and maintenance of conditional SYNGAP1 rescue mouse line (SYNGAP1^ls/+^, a gift from Dr. Gavin Rumbaugh) have been previously described^22^. Inducible CAGG-Cre-ER male mice were purchased from Jackson Laboratories (JAX stock #004682) and crossed to female SynGAP^+/ls^ for genetic rescue experiments. Male and female experimental mice were utilized for all studies in equal numbers and combined when no effect of sex was detected. Age-matched littermates were randomized to treatment groups and a balanced number of SynGAP^+/ls^ and WT mice were used. Mice were group-housed with no more than 5 mice/cage and maintained on a standard 12-h light/12-h dark cycle. Food and water were provided *ad libitum*.

### Genotyping

Crude DNA extract (0.2 μl) (KAPA Biosystems) from ear punch tissue was used for genotyping the SYNGAP1 Lox-stop allele with the following primers: AAGGCTGGGGTTAATTCAGG, CAACCTTCCTACCCTGCTCA, and TTAAGGGCCAGCTCATTCCT (The Jackson Laboratory), or the CRE allele with the following primers: AGGTGGACCTGATCATGGAG (transgene forward), ATACCGGAGATCATGCAAGC (transgene reverse), CTAGGCCACAGAATTGAAAGATCT (internal positive control forward), and GTAGGTGGAAATTCTAGCATCATCC (positive control reverse) using standard polymerase chain reaction protocols.

### Tamoxifen Injections-Cre induction

Tamoxifen (TMX, Sigma T5648, St. Louis, MO) was dissolved in corn oil (Sigma C8267) containing 10% ethanol by sonication to a final TMX dosage of 75 mg/kg, injectable concentration of 10 mg/ml, and volume of 7.5 ml/kg. Starting at PND 21-23, mice were intraperitoneally administered the TMX solution once a day for 3 consecutive days.

### Western blot analysis of full-length SYNGAP1 protein restoration after TMX treatment

All mice used in the genetic rescue experiment were tested for full-length SYNGAP1 protein expression levels by Western blot. Briefly, one hippocampal or cortical section was collected from each animal, snap frozen in liquid nitrogen and stored at −80 °C until further processing. Samples are then homogenized with 12 strokes of a glass-Teflon homogenizer with twisting in 1% deoxycholate lysis buffer with 0.1 M Tris-HCl (pH 9), 10 mM sodium fluoride (Sigma, 201154), 2 mM sodium orthovanadate (Sigma, 450243) and protease/phosphatase inhibitor cocktails (Sigma, P8340/P5726). Following incubation in lysis buffer for 15 min, lysates were centrifuged for 15 min at 13,000 G to remove nuclei and debris. Protein concentration in the supernatant was determined using a Pierce BCA kit (ThermoFisher, 23225). For Western blot, proteins (20 μg per lane) were separated by SDS–polyacrylamide gel electrophoresis and transferred to a polyvinylidene difluoride membrane (Bio-Rad). Membranes were blocked in 4% milk in TBST [0.01 M tris (pH7.4), 0.15 M NaCl, 0.1% Tween 20] for 1 h at room temperature and incubated with primary antibodies overnight at 4 °C. Primary antibodies and dilutions used for Western blots were SYNGAP1 (1:1000, clone D88G1, Cell Signaling, Cat. No. 9479 BF), and β-actin (1:10,000, GT×109639, GeneTex). Primary antibodies were detected using species-specific horseradish peroxidase–conjugated secondary antibodies. Blots were developed using Super Signal West Pico PLUS Substrate (ThermoFisher) and imaged using a Protein Simple imaging system (San Jose, CA). Protein bands were measured by densitometry and quantified as the densitometric signal of SYNGAP1 divided by the densitometric signal of β-actin as loading control.

### Hippocampal slice preparation, stimulation, and lysate preparation

Acute brain slices were prepared from P7, P14 and P60 SYNGAP1^ls/+^ and WT littermate mice using N-methyl-D-glucamine (NMDG) protective recovery method^49^. Mice were deeply anesthetized with isoflurane, brains were removed, and 400 μm-thick coronal hippocampal slices were sectioned in pre-chilled carbogenated NMDG-HEPES Acsf (92 mM NMDG, 2.5 mM KCl, 1.2 mM NaH_2_PO_4_, 30 mM NaHCO_3_, 20 mM HEPES, 25 mM glucose, 2 mM thiourea, 5 mM Na-ascorbate, 3 mM Na-pyruvate, 0.5 mM CaCl_2_.4H_2_O, and 10 mM MgSO_4_.7H_2_O; titrated to pH 7.4 with concentrated hydrochloric acid) using a VT1200s vibrating microtome (Leica). The transcardial perfusion procedure with 30 ml of chilled NMDG-HEPES aCSF was performed on P60 animals prior to brain dissection. Upon completion of the sectioning procedure, each hippocampal slice was placed in an alternate treatment group, with treatment groups being arbitrarily assigned. Slices were collected using a cut-off plastic Pasteur pipet and transferred into a pre-warmed (32-34°C) initial recovery chamber filled with NMDG-HEPES aCSF and incubated for 10-15 min. Slices were then transferred to a carbogenated modified HEPES holding solution (92 mM NaCl, 2.5 mM KCl, 1.2 mM NaH_2_PO_4_, 30 mM NaHCO_3_, 20 mM HEPES, 25 mM glucose, 2 mM thiourea, 5 mM Na-ascorbate, 3 mM Na-pyruvate, 2 mM CaCl_2_.4H_2_O, and 2 mM MgSO_4_.7H_2_O; pH 7.4) for an additional 60–90 min recovery at room temperature. Slices were then incubated in HEPES-aCSF (129 mM NaCl/ 5 mM KCl/ 2 mM CaCl_2_/ 30 mM Glucose/ 25 mM HEPES; pH 7.4) for 1h at 32–34 °C to equilibrate, and subsequently stimulated with NMDA/glycine (100/1 μM), DHPG (100 uM) or HEPES-aCSF control for 5 minutes at 32–34°C. NMDAR agonists NMDA (Sigma, M3262; 100 µM) + glycine (Fisher Scientific, Pittsburgh, PA, USA, BP-381; 1 µM) were dissolved in HEPES-aCSF in the presence of non-NMDA glutamate receptor blockers (CNQX, Tocris, 0190, 40 µM; Nimodipine, Tocris, 0600, 1 µM; LY-367385, Sigma, L4420, 100 µM; MPEP, Tocris, 1212, 10 µM). Type I mGluR agonist DHPG (Tocris, 0805; 100 µM) was dissolved in HEPES-aCSF in the presence of non-mGluR glutamate receptor blockers (CNQX, Tocris, 0190, 40 µM; Nimodipine, Tocris, 0600, 1 µM; D(-)-2-Amino-5 phosphonopentanoic acid (APV), Sigma, A5282, 50 µM). Following treatments, slices were quickly frozen in liquid nitrogen and kept at −80 °C until further processing. Lysates were prepared by homogenization in ice cold 1% NP-40 lysis buffer using 12 strokes of a glass-teflon PYREX homogenizer. Tissue lysates were then transferred to a centrifuge tube, incubated on ice for 15 min, and centrifuged at high speed for 15 min. The protein concentration of the supernatant was determined using a BCA assay (Pierce).

### Cortical brain slicing and lysate preparation

We indended to evoke homeostatic plasticity and/or experience-dependent plasticity by trimming all vibrissae (mystacial whiskers) on one side of a 2-month-old mouse with an electric trimmer while the mouse was under very brief isoflurane anesthesia, as previously described^8^. As we did not detect differences between deprived and non-deprived cortices, the datasets were collapsed into a single group for each genotype. 48-h after whisker deprivation, mice were anesthetized with isoflurane and perfused through the heart with ice-cold 1× phosphate-buffered saline (PBS, w/o Ca^2+^ and Mg^2+^). Brains were immediately removed, placed into cold PBS, and sliced in cold PBS using a VT1200s vibrating microtome (Leica). PBS was saturated with carbogen (95% O2/5% CO2) prior to use to ensure stable pH buffering and adequate oxygenation. Five 500-μm slices through the somatosensory (barrel) cortex were divided into ipsilateral and contralateral hemispheres; the barrel cortex microdissected and kept briefly in ice-cold PBS. All microdissected slices from one hemisphere were then combined, and snap frozen in liquid nitrogen and stored at −80 °C until further processing. Lysates from cortical slices were prepared as for hippocampal slices.

### Quantitative Multiplex co-Immunoprecipitation (QMI)

QMI is an ELISA-like assay that simultaneously measures the abundance (in median fluorescent intensity units or MFI) of ∼400 protein interactions in cell lysaes. In QMI, ∼20 target proteins are simultaneously immunoprecipitated onto dye-encoded flow cytometry beads (one antibody per class of bead), probed with ∼20 fluorophore-coupled antibodies that recognize potentially co-associated proteins, and read on a flow cytometer. QMI was performed as described previously^5,8,9^, see Brown et al^43^ for a detailed video protocol, analysis software code for both R and Matlab, and sample datasets. QMI validation in mouse neurons, including specific validation of each antibody pair, was described previously^5^. Antibody clone names and catalog numbers are the same as in^9^, and are listed in Table S1. A master mix containing equal numbers of each antibody-coupled bead was prepared. Beads were distributed to lysates containing equal amounts of protein (by BCA assay), and incubated overnight on a rotator at 4°C. The next day, beads from each sample were washed twice in cold Fly-P buffer (50 mM tris (pH7.4), 100 mM NaCl, 1% bovine serum albumin, and 0.02% sodium azide) and distributed into twice as many wells of a 96-well plate as there were probe antibodies (for technical duplicates). Biotinylated detection (probe) antibodies were added to the wells and incubated at 4°C with gentle agitation for 1 hour. The resulting bead-probe complexes were washed three times with Fly-P buffer, incubated for 30 min with streptavidin-phycoerythrin at 4°C with gentle agitation, washed another three times, resuspended in 125 μl of ice-cold Fly-P buffer, and read on a customized refrigerated Bio-Plex 200 (as detailed in ref^43^).

### Quantification and statistical analysis of QMI data

#### Data preprocessing and inclusion criteria

For each well from a data acquisition plate, data were processed using the BioPlex200’s built-in software to (i) eliminate doublets on the basis of the doublet discriminator intensity (> 5000 and < 25 000 arbitrary units; Bio-Plex 200), (ii) identify the bead class of each bead, which indicates the IP antibody on the bead, and (iii) identify the bead phycoerythrin fluorescence measurement, which indicates the amount of co-associated probe. This process generates a distribution of fluorescence intensity values for each pairwise protein interaction measurement. XML output files were parsed to acquire the raw data for use in MATLAB using the ANC program^42^. No specific analysis was performed on the data to test for outliers.

#### Adaptive non-parametric analysis with empirical alpha cutoff (ANC)

ANC is used to identify high-confidence, statistically significant differences (corrected for multiple comparisons) in bead distributions for each pairwise protein interaction. ANC analysis was conducted in MATLAB (version 2013a) as described in^42^. ANC requires that hits be present in > 70% of experiments at an adjusted p < 0.05. The a-cutoff value required per experiment to determine statistical significance was calculated to maintain an overall type I error of 0.05 (adjusted for multiple comparisons with Bonferroni correction), with further empirical adjustments to account for technical variation. No assessment of normality was carried out as ANC analysis is a non-parametric test.

#### Correlation Network Analysis (CNA)

Modules of protein-protein interactions that co-varied with experimental conditions were identified using CNA as described in ^42,43^. Briefly, bead distributions used in ANC were collapsed into a single median fluorescent intensity (MFI) for each interaction and averaged across technical replicates for input into the WGCNA package for R^44^. Interactions with MFI < 100 were removed as noise, and batch effects were corrected using COMBAT^50^ with ‘experiment number’ as the batch. Power values giving the approximation of scale-free topology were determined using soft thresholding with a power adjacency function. Modules whose eigenvectors significantly correlated with an experimental trait (p < 0.05) were considered ‘‘of interest.’’ Interactions belonging to a module of interest and whose probability of module membership in that module was < 0.05 were considered significantly correlated with that trait.

#### ANCՈCNA

Interactions that were significant by both ANC and CNA for a given experimental condition were considered significantly altered in that condition. Only these ‘double-significant’ interactions are reported in the text and figures.

#### Data visualization

For heatmap visualization of MFI values of CNA module members, log2 transformed data were input into the Heatmap.2 program in R studio, which normalized the data by row for visualization of multiple analytes spanning a 3-log range. All interactions visualized were ANCՈCNA significant for at least one condition. The number of biological replicates for each experiment can be found in figure legends, error bars represent SEM.

## Author contributions

VS performed all experiments with assistance from FH and BK. VS and SEPS analyzed the data. VS and SEPS wrote the manuscript with input from all authors. SEPS initiated the project and acquired funding. GR provided SynGAP^+/ls^ mice and contributed critical scientific discussion and manuscript edits. All authors read and approved the manuscript.

## Acknowledgments

We wish to thank members of the SEPS lab, the Center for Integrative Biology, particularly Damon Page, Kathleen Millen and Brock Grill for helpful discussions and advice. Funding was provided by the National Institute of Mental Health R01 MH113545 (to SEPS), the National Institute of Neurological Disorders and Stroke NS143924 (to SEPS), a FRAXA foundation fellowship (to VS), and financial support from Seattle Children’s Research Institute.

## Conflicts of Interest

The authors declare that they have no competing financial interests.

## Notes

### Competing Interest Statement

The authors have declared no competing interest.

