## Supplementary Material for "Genetic rescue of disrupted synaptic protein interaction network dynamics following *SYNGAP1* reactivation"

**Affiliations:**

**Keywords:**

**Contents:**

Table S1

Figures S1-S3

### Table S1. IP probe antibody panels used for QMI

#### Scaffold and structural proteins

| Target | QMI Component | Clone | Supplier | Cat# | RRID |
|---|---|---|---|---|---|
| Homer1 | IP<br>Probe | AT1F3<br>D3 | LifeSpan Bioscience<br>Santa Cruz | LS-C103482<br>sc-17842 | AB_2264378<br>AB_627742 |
| NL3 | IP<br>Probe | 566209<br>G2 | R&D Systems<br>Santa Cruz | MAB6088<br>sc-271880 | AB_2044707<br>AB_10709173 |
| panShank | IP<br>Probe | NA<br>N23B/49 | NA<br>Neuromab | NA<br>75-089 | NA<br>AB_10672418 |
| PSD-95 | IP<br>Probe | K28/74<br>K28/43 | Biolegend<br>Biolegend | 810301<br>810401 | AB_2564749<br>AB_2564750 |
| SAPAP | IP<br>Probe | NA<br>N127/31 | NA<br>Biolegend | NA<br>832601 | NA<br>AB_2564958 |
| SAP97 | IP<br>Probe | RPI197.4<br>2D11 | Enzo Life Sciences<br>Santa Cruz | ADI-VAM-PS005<br>sc-9961 | AB_10618588<br>AB_2092015 |
| Shank1 | IP<br>Probe | N22/21<br>N22/21 | Neuromab<br>Neuromab | 75-064<br>75-064 | AB_2270283<br>AB_2270283 |
| Shank3 | IP<br>Probe | N367/62<br>N69/46 | Neuromab<br>Neuromab | 75-344<br>75-109 | AB_2315921<br>AB_2187730 |

#### Receptor Proteins

| Target | QMI Component | Clone | Supplier | Cat# | RRID |
|---|---|---|---|---|---|
| GluR1 | IP<br>Probe | poly1504<br>N355/1 | Millipore<br>Biolegend | AB1504<br>819801 | AB_2113602<br>AB_2564834 |
| GluR2 | IP<br>Probe | L21/32<br>Poly | Biolegend<br>ProteinTech | 810501<br>11994-1-AP | AB_2564751<br>AB_2113725 |
| mGluR5 | IP<br>Probe | 5675<br>N75/3 | Millipore<br>Neuromab | AB5675<br>75-115 | AB_2295173<br>AB_10672260 |
| NMDAR1 | IP<br>Probe | 54.1<br>N/A | ThermoFisher<br>N/A | 32-0500<br>N/A | AB_2533060<br>N/A |
| NMDAR2A | IP<br>Probe | N327/95<br>Poly | Biolegend<br>LifeSpan Bioscience | 832101<br>LS-C166676 | AB_2564953<br>AB_3717840 (pending) |
| NMDAR2B | IP<br>Probe | N59/20<br>N59/36 | Biolegend<br>Biolegend | 832501<br>818701 | AB_2564957<br>AB_2564823 |

#### Signaling proteins

| Target | QMI Component | Clone | Supplier | Cat# | RRID |
|---|---|---|---|---|---|
| CaMKII | IP<br>Probe | 6G9<br>polyC6970 | Enzo<br>Sigma | ADI-KAM-CA002<br>C6974 | AB_10617228<br>AB_258984 |
| Fyn | IP<br>Probe | Fyn59<br>Fyn15 | Biolegend<br>Santa Cruz | 626502<br>sc-434 | AB_2278824<br>AB_627642 |
| PI3K p85 | IP<br>Probe | U5<br>AB6 | ThermoFisher<br>Millipore | MA1-74183<br>05-212 | AB_2163452<br>AB_309658 |
| PIKE | IP<br>Probe | 263A<br>DN8 | Bethyl Laboratories<br>Rockland | A304-263A<br>200-401-DN8 | AB_2620459<br>AB_2612161 |
| SynGap | IP<br>Probe | D20C7<br>D78B11 | Cell Signaling<br>Cell Signaling | 5539<br>5540 | AB_10694401<br>AB_10695900 |
| Ube3a | IP<br>Probe | E4<br>Poly | Santa Cruz<br>Bethyl Labs | sc-166689<br>A300-352A | AB_2211807<br>AB_185564 |

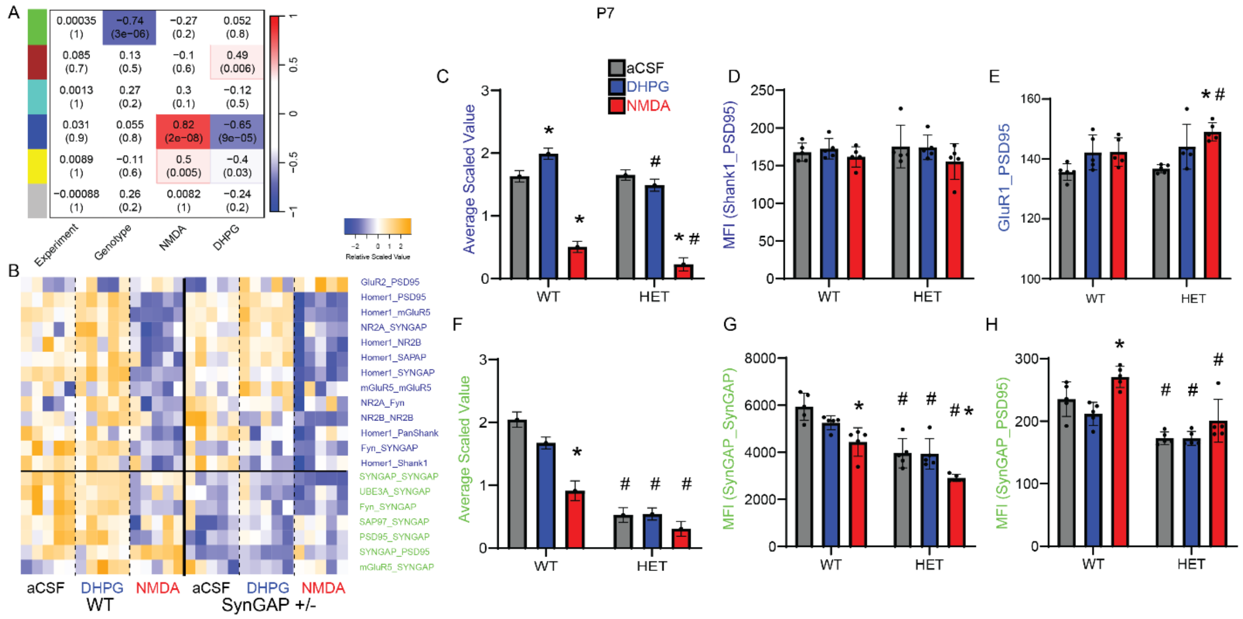


**Figure S1: Protein interaction network response to NMDA in P7 SynGAP$^{+/ls}$ hippocampal slices.** A) A module-trait table shows the correlation coeficent (top number) and p-value (bottom number) for the correlation between each module's eigenvector and experimental variables listed at bottom. B) A row-scaled heatmap shows all interactions that were both significant in binary comparisons by adaptive nonparametric test corrected for multiple comparisons (ANC) and a member of a CNA module. Interactions listed at right are colored by module membership. C-H) Bar graphs represent the average scaled value (G,F) and example interactions (D-E, G-H) for the blue NMDA-correlated (C-E) or the green SynGAP-correlated (F-H) modules. * indicates p<0.05 comparing aCSF vs treatment within-genotype; # indicates p<0.05 comparing WT vs. SynGAP$^{+/ls}$ within-treatment, by 2-way ANOVA followed by Tukey's post-hoc testing.

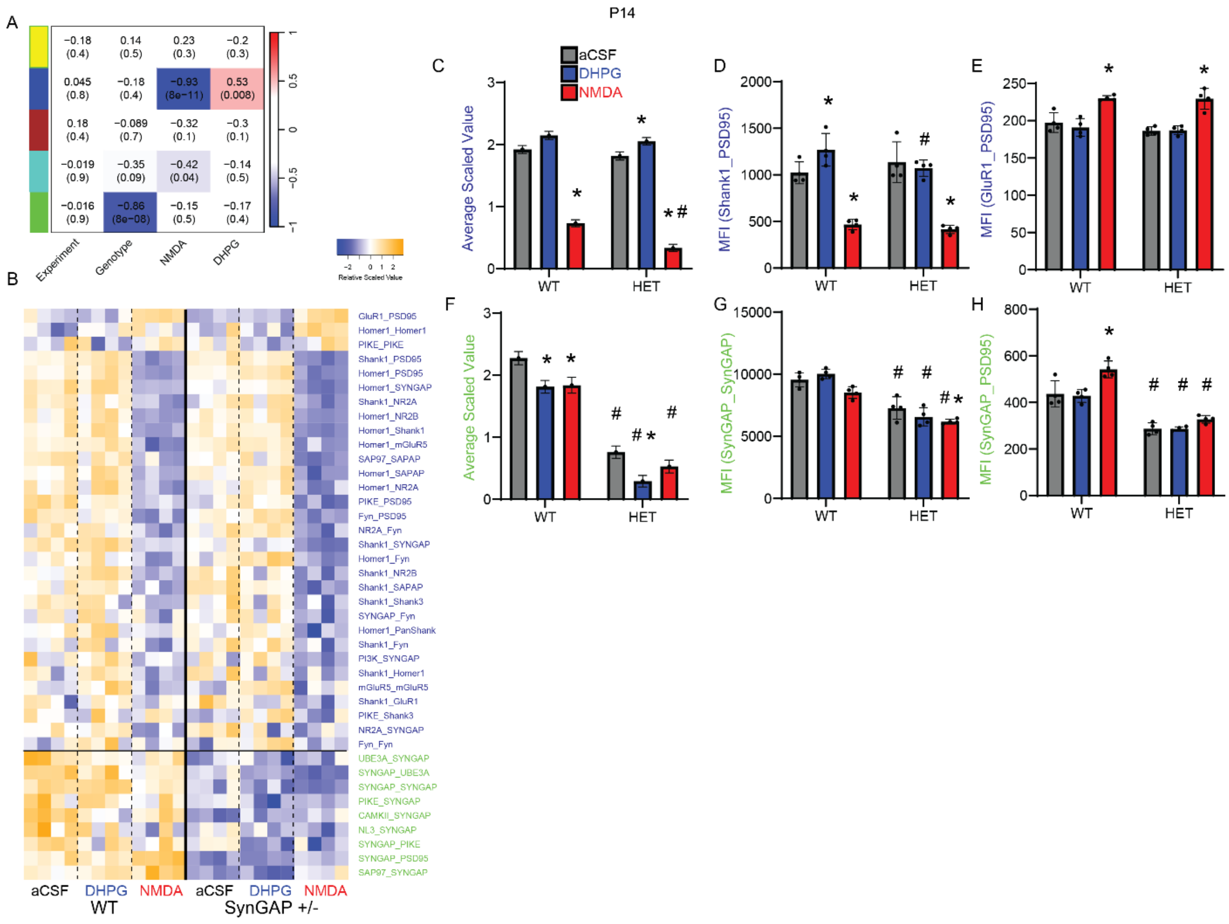


**Figure S2: Protein interaction network response to NMDA in P14 SynGAP$^{+/ls}$ hippocampal slices.** A) A module-trait table shows the correlation coeficent (top number) and p-value (bottom number) for the correlation between each module's eigenvector and experimental variables listed at bottom. B) A row-scaled heatmap shows all interactions that were both significant in binary comparisons by adaptive nonparametric test corrected for multiple comparisons (ANC) and a member of a CNA module. Interactions listed at right are colored by module membership. C-H) Bar graphs represent the average scaled value (G,F) and example interactions (D-E, G-H) for the blue NMDA-correlated (C-E) or the green SynGAP-correlated (F-H) modules. * indicates p<0.05 comparing aCSF vs treatment within-genotype; # indicates p<0.05 comparing

WT vs. SynGAP$^{+/ls}$ within-treatment, by 2-way ANOVA followed by Tukey's post-hoc testing.

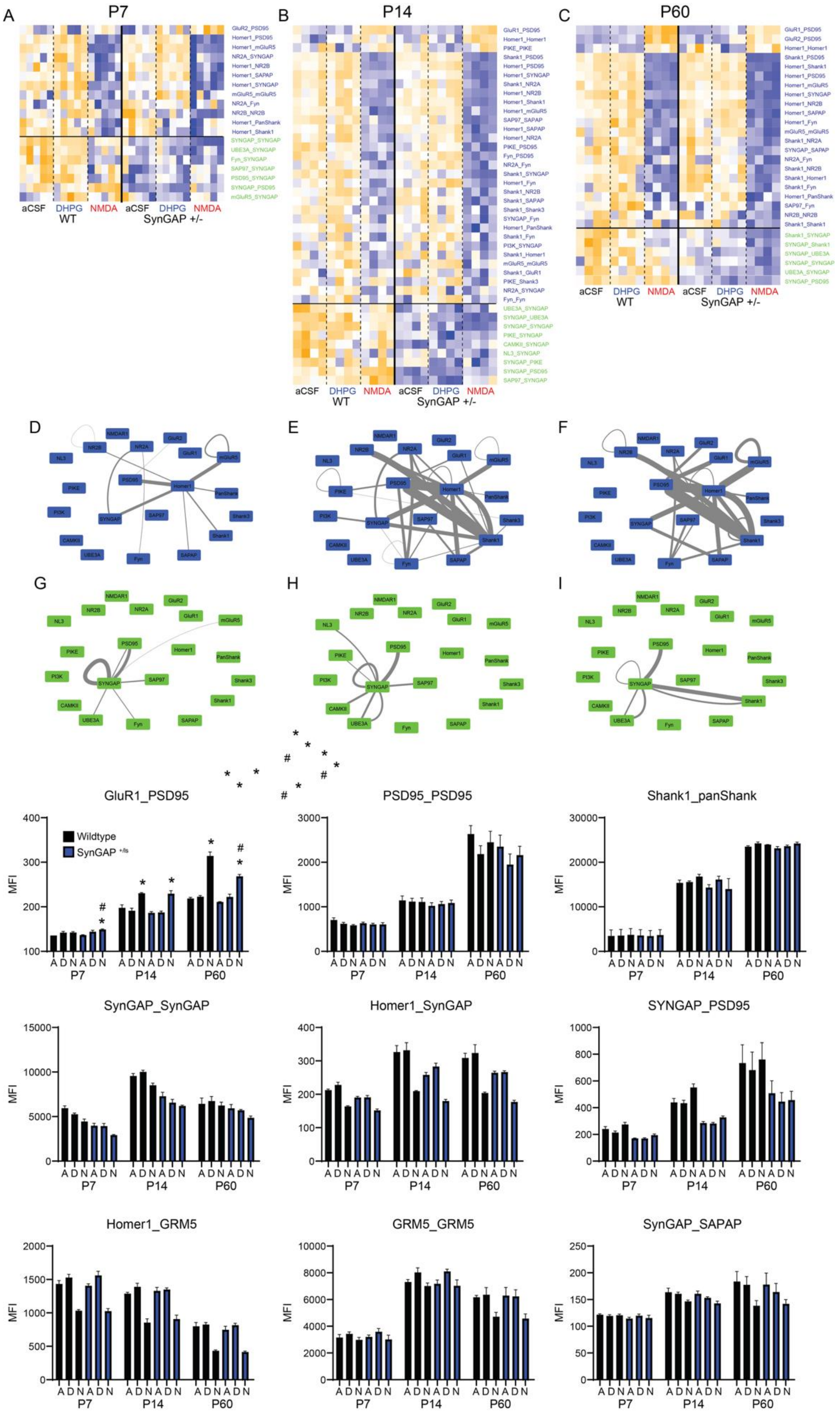
A
P7
aCSF DHPG NMDA aCSF DHPG NMDA
WT SynGAP +/-
B
P14
aCSF DHPG NMDA aCSF DHPG NMDA
WT SynGAP +/-
C
P60
aCSF DHPG NMDA aCSF DHPG NMDA
WT SynGAP +/-
D
E
F
G
H
I
GluR1_PSD95
Wildtype
SynGAP +/fs
PSD95_PSD95
Shank1_panShank
SynGAP_SynGAP
Homer1_SynGAP
SYNGAP_PSD95
Homer1_GRM5
GRM5_GRM5
SynGAP_SAPAP
MFI
A D N A D N A D N A D N A D N A D N
P7 P14 P60

**Figure S3: Comparison of protein interaction network response to NMDA across development in SynGAP$^{+/ls}$ hippocampal slices.** A-C) Row-scaled heatmaps for each age (P7, P14, P60) show all interactions that were both significant in binary comparisons by adaptive nonparametric test corrected for multiple comparisons (ANC) and a member of a CNA module. Interactions listed at right are colored by module membership. These heatmaps are identaical to Fig 1, S1 and S2 and are re-displayed for direct comparison. D-I) Node-edge diagrams show interactions in the NMDA (D-F) and SynGAP (G-I) modules for P7 (D,G), P14 (E,H) and P60 (F,I). Edge width indicates the magnitude of the fold-change for NMDA vs aCSF or WT vs SynGAP$^{+/ls}$. J-R) Bar graphs show the median fluorescent values for example interactions across ages. * indicates $p<0.05$ comparing aCSF vs treatment within-genotype; # indicates $p<0.05$ comparing WT vs. SynGAP$^{+/ls}$ within-treatment, by 2-way ANOVA followed by Tukey's post-hoc testing. All comparisons are within-age groups. Experimental design and normalization strategy does not allow for direct statistical comparisons across age groups.
